# Hybrid short- and long-read assembly of an experimentally evolved *Sodalis glossinidius* strain SgGmmC1*

**DOI:** 10.1101/2025.06.26.659281

**Authors:** Poppy Pescod, Lee R. Haines, Alistair C Darby, Ian Goodhead

**Author notes:** Address correspondence to Ian Goodhead. These authors contributed equally to this work (Poppy Pescod, Lee R. Haines).

## Abstract

Bacterial symbionts of insects undergo dramatic genome reduction during their evolutionary transition from free-living to host-dependent lifestyles, but the dynamics of this process remain poorly understood due to the difficulty of observing these processes in real-time. *Sodalis glossinidius*, a facultative endosymbiont of tsetse flies, provides an exceptional opportunity to study this transition experimentally: Unlike highly specialised obligate symbionts, *S. glossinidius* can be cultured *in vitro* and retains a large genome (4 Mbp) with extensive pseudogene content (49%), suggesting a recent evolutionary transition. Here, we present a comparative genomic analysis of *S. glossinidius* strains isolated from laboratory colony-derived *Glossina morsitans morsitans*, comparing one after ten years of serial passaging in laboratory culture (SgGmmC1*) to a counterpart isolated at the same time (SgGmmB4). Hybrid genome assembly using Oxford Nanopore and Illumina technologies produced a high-quality 4.29 Mbp genome comprising one circular chromosome and four plasmids similar. Comparative analysis revealed a significant deletion (17,209 bp) containing 31 genes, including (involved in thiamine biosynthesis) and genes encoding sulfur transporters. Additionally, we identified multiple small-scale mutations (8 deletions, 39 insertions, 10 SNPs) resulting in frameshifts in genes including a hemolysin precursor (*shlA*). Our findings demonstrate that, under stable laboratory conditions without the selective pressures of the host environment, *S. glossinidius* continues to undergo genome degradation. The loss of *thiM* supports previous hypotheses of complementary metabolic pathways between *S. glossinidius* and the primary symbiont *Wigglesworthia glossinidia* for thiamine biosynthesis. This study provides insights into the evolutionary trajectory of facultative symbionts and has implications for paratransgenic approaches using *S. glossinidius* for trypanosome control.

**DATA SUMMARY:** Raw Nanopore and MiSeq reads are available in the European Nucleotide Archive under Project PRJEB32321 (Read accessions: ERX3321202 and ERX3321201 respectively). A full GBK formatted annotation of SgGmmC1* has been deposited in Figshare at https://doi.org/10.17866/rd.salford.8052437.v1.

**IMPACT STATEMENT:** In this study we provide new insights into the ongoing genome degradation of *Sodalis glossinidius*, a facultative endosymbiont of tsetse flies implicated in their ability to transmit trypanosomiasis, during prolonged laboratory culture. Through high-quality comparative genomics we reveal significant gene loss and mutational changes, including disruption of pathways involved in nutrient biosynthesis. These findings enhance our understanding of symbiont genome evolution and inform the development of *S. glossinidius* as a potential tool for paratransgenic strategies in controlling trypanosome transmission.

## INTRODUCTION

Tsetse flies (*Genus: Glossina*) are unusual insects that, like keds and bat flies, reproduce by giving birth to live young (adenotrophic viviparity). As obligate haematophages, tsetse harbours a limited bacterial microbiome due to their restricted diet, feeding solely on blood in the adult life stage and on proteinaceous milk from the female’s specialised accessory glands during the larval stage (Minchin, 1905). At least four species of bacteria play important roles in the nutrition, fecundity and vectorial capacity of tsetse flies: Wigglesworthia *glossinidia*, an obligate primary symbiont is ubiquitous; *Sodalis glossinidius*, a facultative commensal symbiont, and *Wolbachia* and *Spiroplasma*, parasitic reproductive manipulators, all have varied infection rates across the wide distribution of tsetse across sub-Saharan Africa (Doudoumis *et al*, 2017). *S. glossinidius* is a Gram-negative secondary bacterial endosymbiont that has been linked to modifying tsetse fly susceptibility to trypanosome infection by parasites belonging to the genus *Trypanosoma* (Trappeniers *et al*, 2019; Makhulu *et al*, 2021; Kallu *et al*, 2023). Trypanosomes cause a fatal disease in humans and animals called trypanosomiasis (Maudlin and Ellis, 1985).

Obligate insect endosymbiont genomes demonstrate specialisation to their host environment by being non culture-viable and having small, compact genomes relative to free-living relatives (Moran *et al*., 2008). *S. glossinidius* is notable for being the first endosymbiont discovered to be both amenable to *in vitro* culture (Dale & Maudlin, 1999) and for having a relatively large genome: The *S. glossinidius* genome is 4 Mbp, similar in size to free-living bacteria, but large in comparison to the 0.7 Mbp genome of *W. glossinidia*; both of these features suggest a recent switch to host association and symbiosis (Toh *et al*., 2006).

The transition of bacterial genomes from free-living to endosymbiosis is poorly understood. Evidence suggests endosymbiont evolution involves inactivation of functional genes via small insertions or deletions to produce “pseudogenes”. These pseudogenes are subsequently lost in large deletion events facilitated by frequent population bottlenecks (Goodhead and Darby, 2015). As mutations accumulate in non-essential genes, frameshifts and large numbers of insertions or deletions (indels) result in loss of gene function in a process known as pseudogenisation; pseudogenes may or may not retain function depending on the level of degradation (Goodhead *et al*, 2020).

*S. glossinidius* represents a unique mid-way point in the evolutionary transition towards obligate endosymbiosis, evidenced by the high proportion of pseudogenes in its genome: 49% compared to the usual 1% present in free-living bacteria (Goodhead *et al*., 2017; Toh *et al*., 2006). However, whilst the obligate endosymbiont *W. glossinidia* is present in all individuals of the Glossina genus, *S. glossinidius* is found at rates ranging from 1% - 94% in wild tsetse populations (*Farikou et al* 2011; Dennis *et al*, 2014; Mfopit *et al*, 2023). This lack of consistency implies *S. glossinidius* retains a facultative relationship with its host and impacts our ability to study bacterial genome degradation in natural settings. Indeed, genomic information for *S. glossinidius* has largely been collected from isolates derived from *Glossina* under laboratory conditions (Toh *et al*., 2006; Belda *et al*., 2010; Goodhead *et al*., 2017).

*S. glossinidius* has several attributes that make it a promising microorganism for use in tsetse and trypanosome control strategies – it is amenable to culture *in vitro*, has a demonstrable ability to infect the target vector, and can be genetically manipulated to express control factors (Beard *et al*, 1993; Welburn *et al*, 1987). A recombinant strain of *S. glossinidius* expressing functional anti-trypanosome Nanobodies^®^ has been used to deliver the Nanobodies to tsetse in an *in vivo* lab-based proof of concept study (De Vooght *et al* 2012). Whilst promising, the effector molecule was only stable within the tsetse host when its native *S. glossinidius* population was significantly suppressed. This suggests fitness differences between native and lab-reared strains of the bacterium, meaning further studies into the fitness of lab-derived *S. glossinidius* versus natural populations are warranted. A thorough understanding of the pathway of genome degradation in lab-based systems will also be vital for paratransgenic approaches, to predict impacts on bacterial fitness or their ability to successfully infect the tsetse host.

Here we present a high-quality whole genome sequence assembly and annotation of a strain of *S. glossinidius* isolated from a lab colony of *Glossina morsitans morsitans* that was serially passaged *in vitro* for ten years. We have compared its genome to that of a counterpart isolated at the same time and from the same colony as the original strain, with the aim of directly observing genome reduction under laboratory settings. We identify both small and large-scale deletions as well as smaller scale mutations, produce a better understanding of pathways under selection in laboratory settings, and hypothesis the impact on natural *S. glossinidius* populations.

## METHODS

### In vitro culture of *S. glossinidius*

*S. glossinidius* was isolated from two male *G. m. morsitans* from the colony at the Liverpool School of Tropical Medicine in 2007 following previously described methods (Matthew *et al*., 2005). The strains were named SgGmmC1 and SgGmmB4, and were immediately stored at −80°C. The SgGmmB4 isolate was sequenced following a single passage from a glycerol stock as previously described using Pacific Biosciences RSII sequencing (Goodhead *et al*, 2020). Beginning in 2009, cultures of SgGmmC1 were incubated at 27°C in a microaerophilic atmosphere and passaged every 10-14 days for ten years, alternating between solid-phase growth on Columbia agar base supplemented with defibrinated horse blood (Oxoid/Thermo Fisher) and liquid-phase growth in serum-free insect media (Sigma Aldrich). We here refer to the “evolved” strain of *Sodalis glossinidius* as “SgGmmC1*”.

### Comparative genome analysis

DNA was extracted from a cell pellet of SgGmmC1* using the Zymo Quick DNA kit and quantified using a Qubit™ 3.0. The SgGmmC1* genome was sequenced using both Oxford Nanopore MinION and Illumina MiSeq platforms. A library for the MinION was generated using the SQK-LSK108 1D ligation kit and sequenced on a R9.4 flowcell, producing 149,847 reads of mean length 3,477.62bp and length N50 of 14,468bp. A total of 521,111,234 bases were sequenced giving an estimated 118x genome coverage. For Illumina MiSeq sequencing, a QIAseq FX DNA library prep kit and v2 150bp sequencing cartridge were used producing a total of 755,405 paired-end reads. MiSeq reads were assessed and adaptors trimmed using FastQC (v0.11.4) (Andrews, 2010). MiSeq reads were trimmed for quality using fastp with default settings (v0.12.5) (Chen *et al*., 2018).

A hybrid genome for SgGmmC1* was built using Unicycler (v0.4.8-beta) (Wick *et al*., 2017) by assembling the MiSeq reads *de novo* and mapping the long MinION reads onto this assembly for structural correction. The hybrid assembly was annotated using PROKKA (v1.13.3) (Seemann, 2014) using the reference strain SgGmmB4 genome (GenBank Accession LN854557.1), which was deposited in 2016. The annotated SgGmmC1* assembly was compared to SgGmmB4 using the Artemis Comparison Tool (ACT) (v13.0.0) (Carver *et al*., 2005) to identify structural changes and large deletions, and Snippy (v3.1) (Seemann, 2017) to identify single-nucleotide polymorphisms (SNPs) and insertions/deletions. An assembly graph of the assembly was produced using Bandage (v0.8.1) (Wick *et al*., 2015) to assess its quality (Data not shown).

## RESULTS

### Comparative genomics of SgGmmC1* and SgGmmB4

The assembled SgGmmC1* genome was 4,288,975bp in size with 54.43% GC and no unidentified base pairs (Ns). The assembly consisted of six contigs: one circular chromosome (4,149,326bp), four circular plasmids (pSG1, 81,522bp; pSG2, 26,425bp; pSG3, 19,512bp; and pSG4/3, 10,816bp) and one plasmid fragment (pSG2 plasmid frag, 1,374bp) were produced. SgGmmC1* was identified with 99.99% similarity to the reference strain SgGmmB4 using BLASTN (v2.8.1) (Figure 1) (Zhang *et al*., 2000). One large deletion of 17, 209bp was identified, containing 13 coding genes and 18 non-coding genes (Table 1). This deleted region included the coding gene *thiM*, which encodes a protein involved in thiamine biosynthesis and one of two copies of esar that encodes a protein related to the quorum-sensing protein LuxR. Further deleted genes included three sulfur and one sulfite transmembrane transporter proteins; one copy of the peptidase yxeP (the other copy of which is pseudogenised elsewhere in the genome); and multiple sugar phosphotransferases (Figure 1 and Table 1).

**Table 1.**
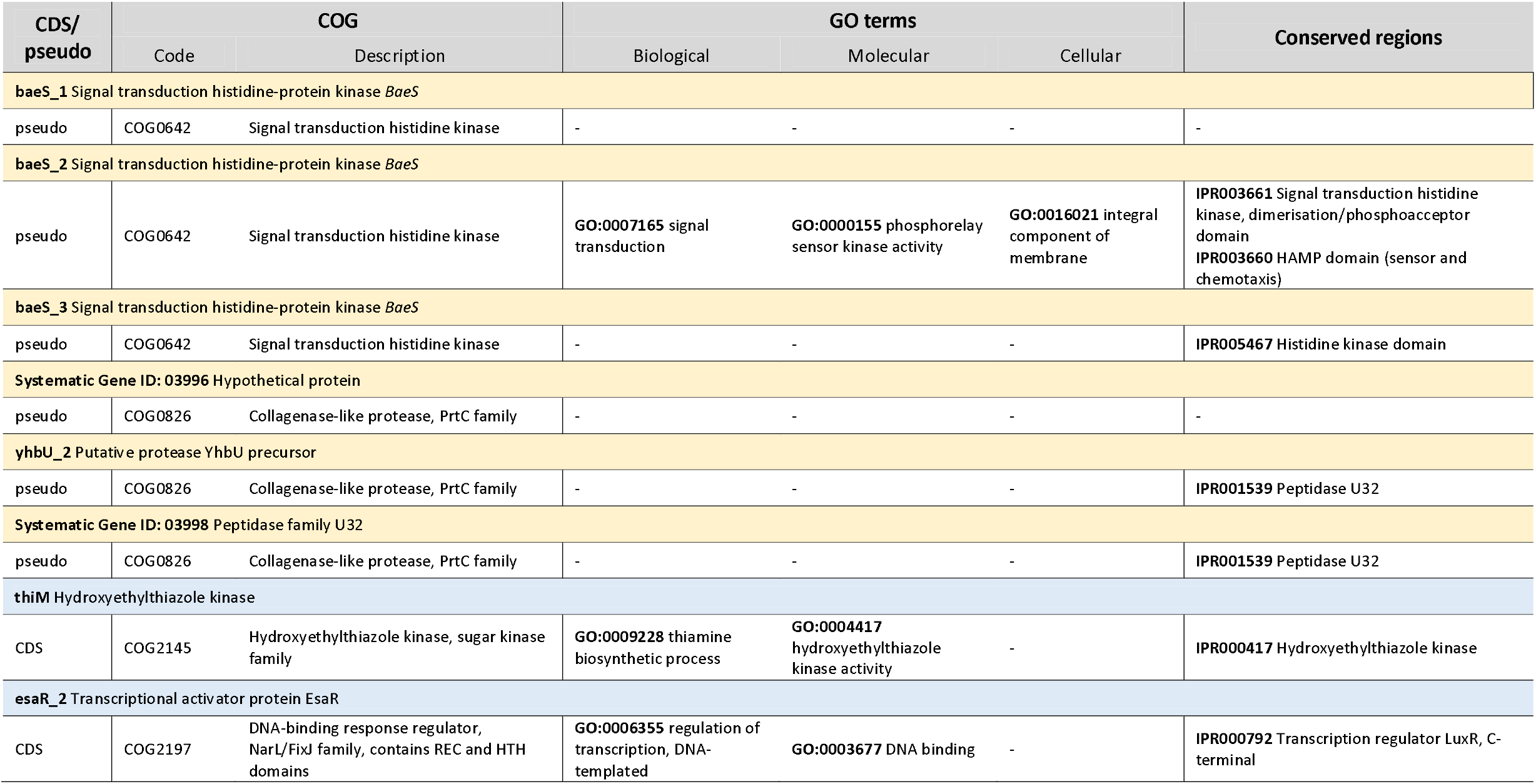
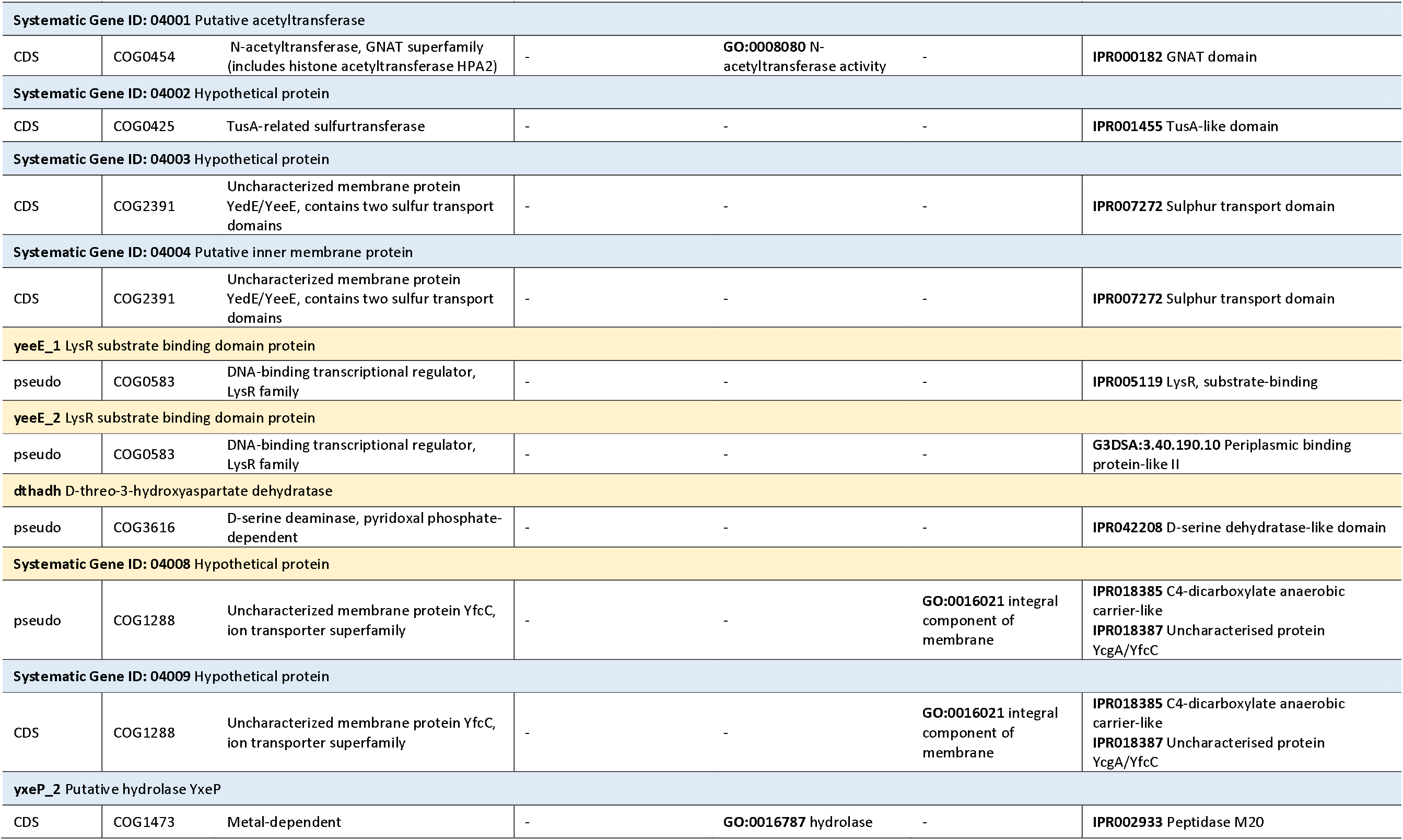
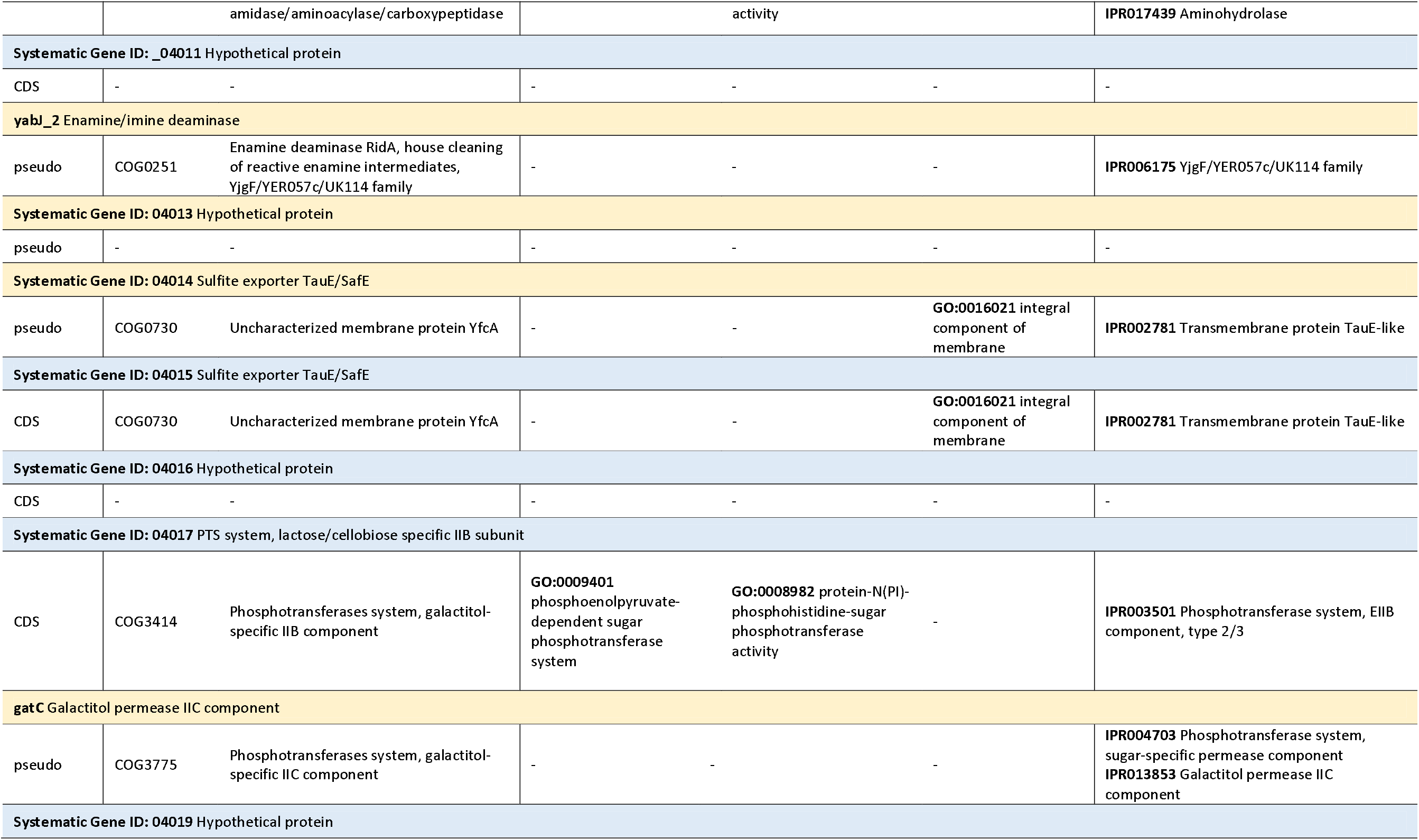
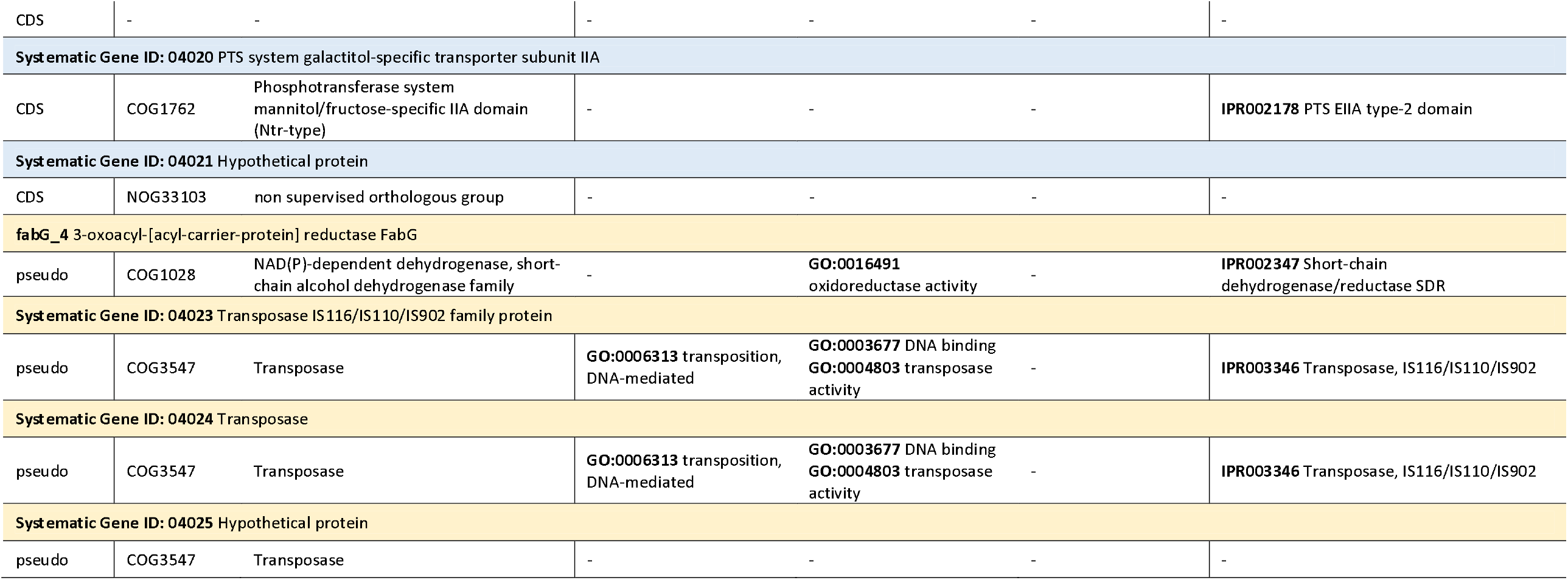
Genes within the 17,209 bp deleted region of *Sg*GmmC1* according to the SgGmmB4 reference (Goodhead *et al*, 2020). Gene names and products were annotated using PROKKA; conserved orthologous groups were assigned using the STRING database; gene ontology and conserved regions were assigned using the Interpro database. Key: CDS – coding DNA sequence; pseudo – pseudogene; COG – clustered orthologous group; GO – gene ontology. Yellow – pseudogenes; blue – CDS.

**Figure 1:**
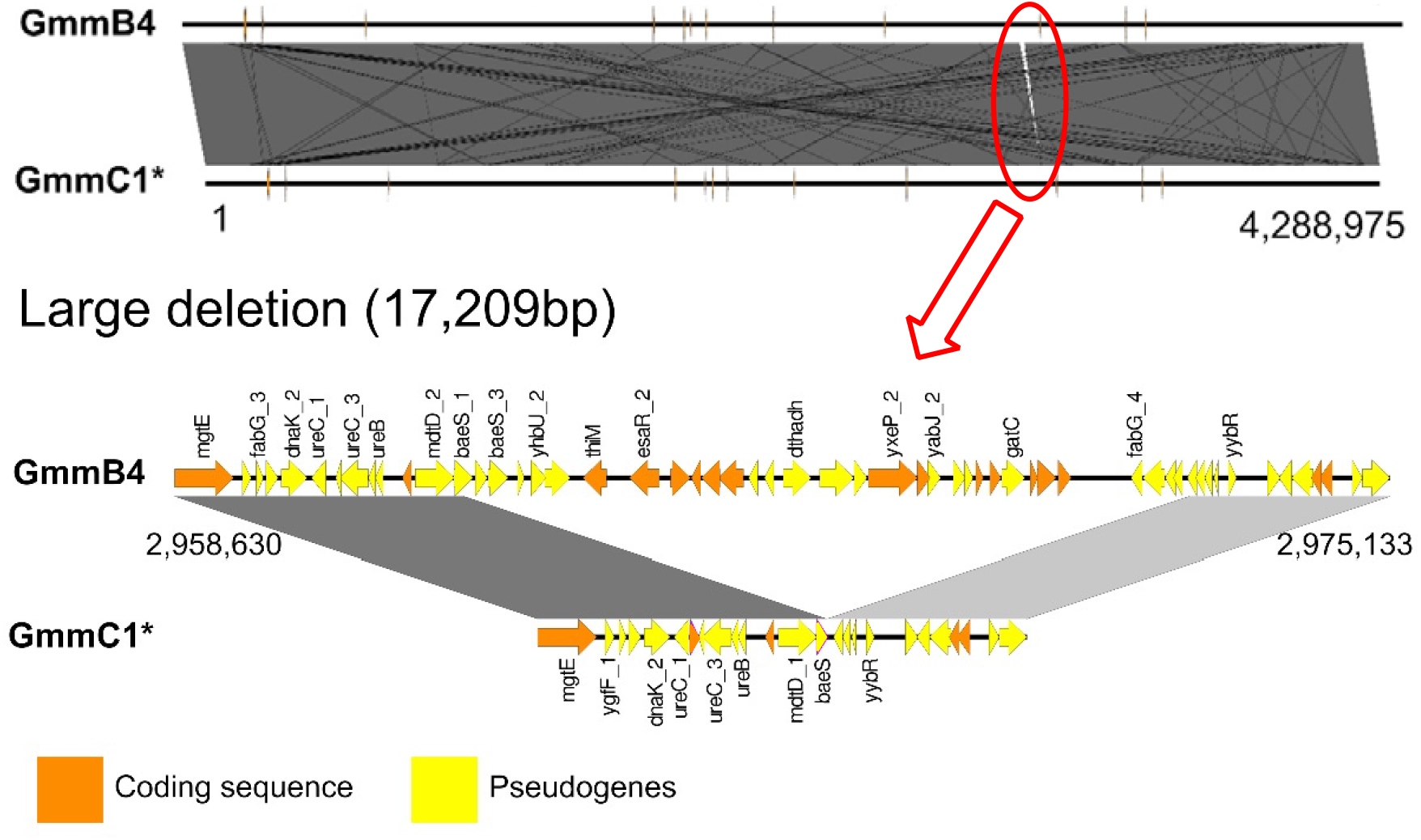
Large deletion in the SgGmmC1* genome. **Top:** Full genome alignment of *Sodalis glossinidius* evolved strain SgGmmC1* against ancestor strain SgGmmB4 showing 99.99% similarity, with the single large deletion expanded to show details of the genes deleted. The top figure shows the alignment of the entire chromosome of each strain, showing the high level of similarity between the sequencies, with the large deletion circled. The bottom diagram shows the coding genes (orange) and pseudogenes (yellow) flanking and included in the deletion. Genome positions (bp) are relative to the SgGmmB4 reference genome. Figures were produced using easyfig.

Snippy also identified eight small deletions (range of 1-27bp, mean 11.6bp, standard deviation 6.08), 39 insertions (1-8bp, mean 3.1bp, std 4.72) and 10 SNPs. In total, 12/32 (37.5%) insertions in a coding region resulted in a frameshift; Putatively pseudogenised genes included the putative transport protein gene *HsrA*, one of five copies of the proline transporter gene *prop* and a hemolysin precursor *shlA* (Table 2).

**Table 2.**
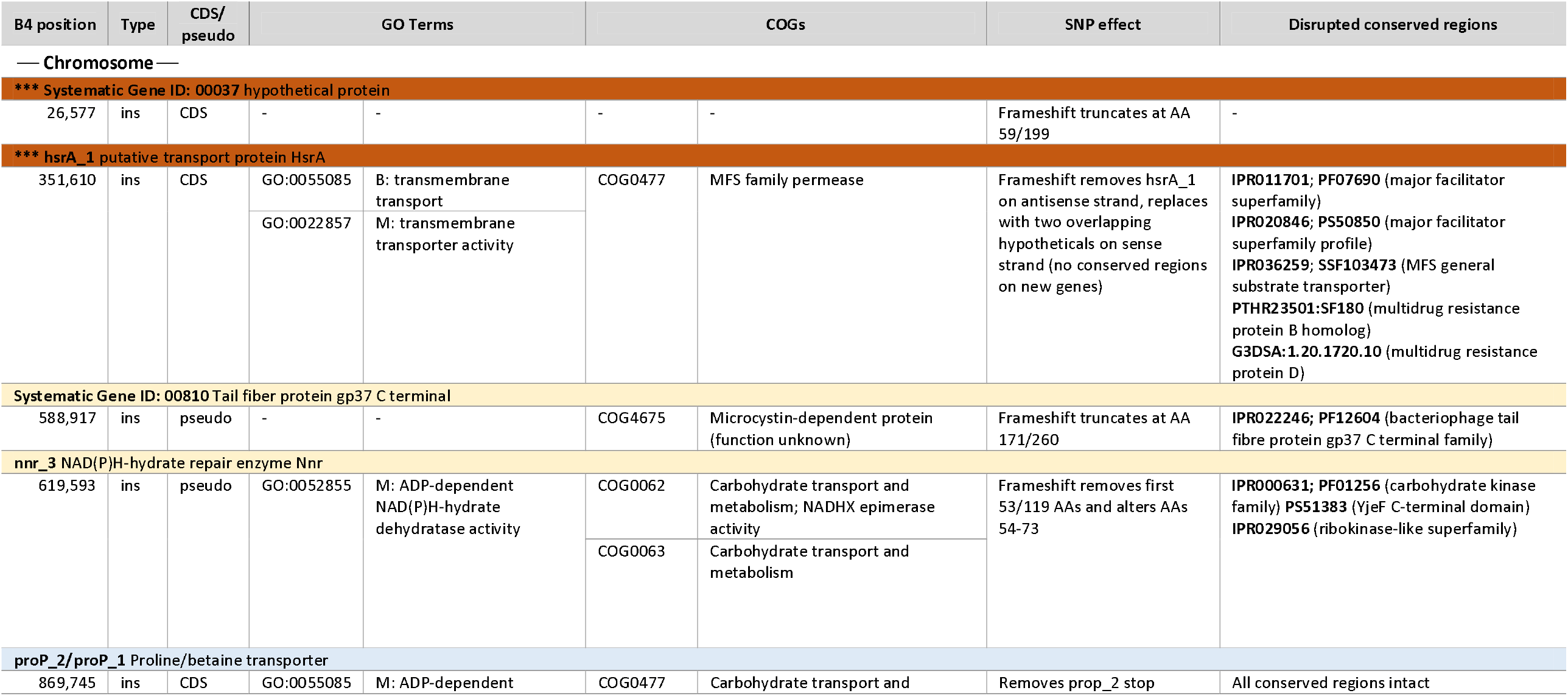
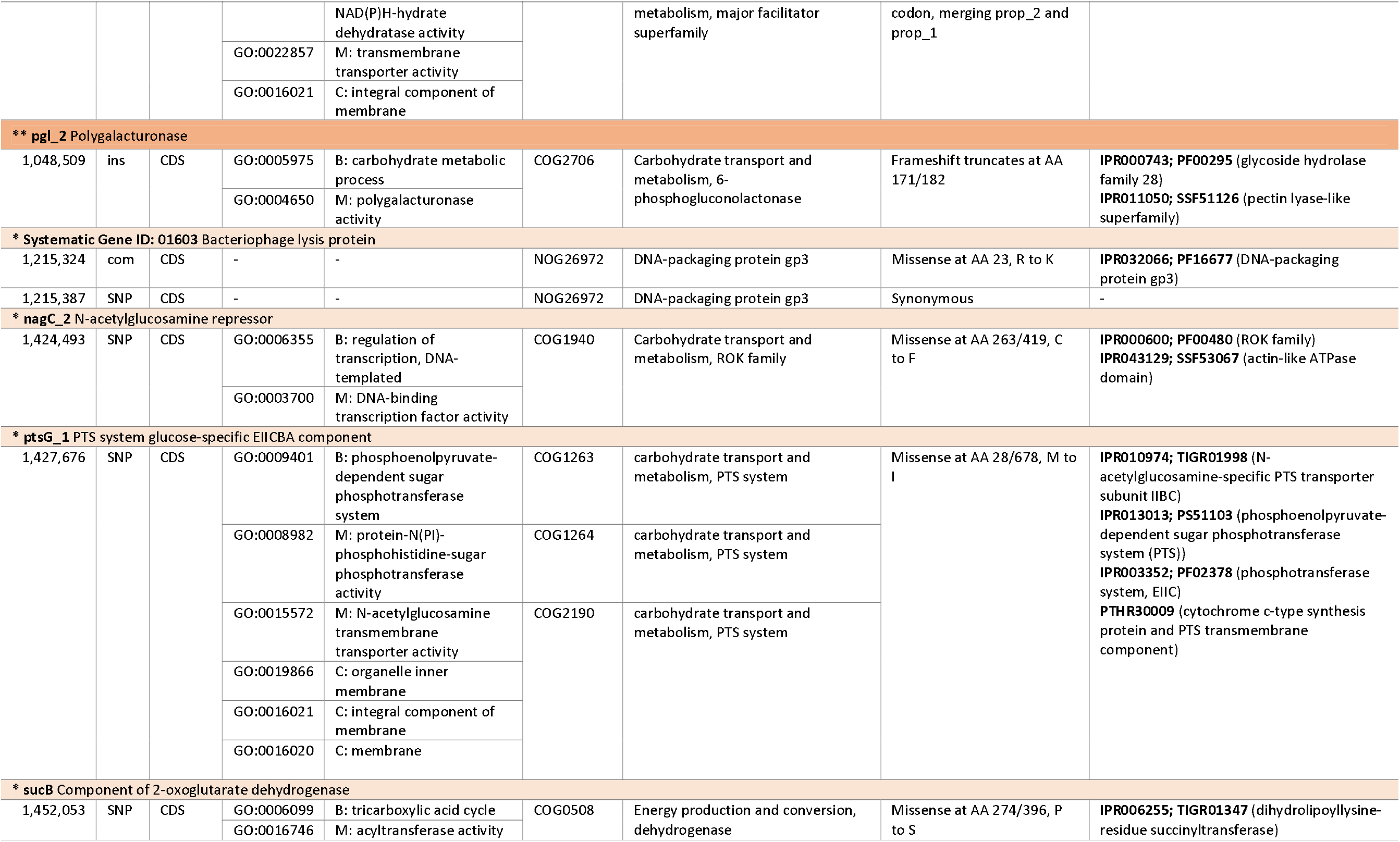
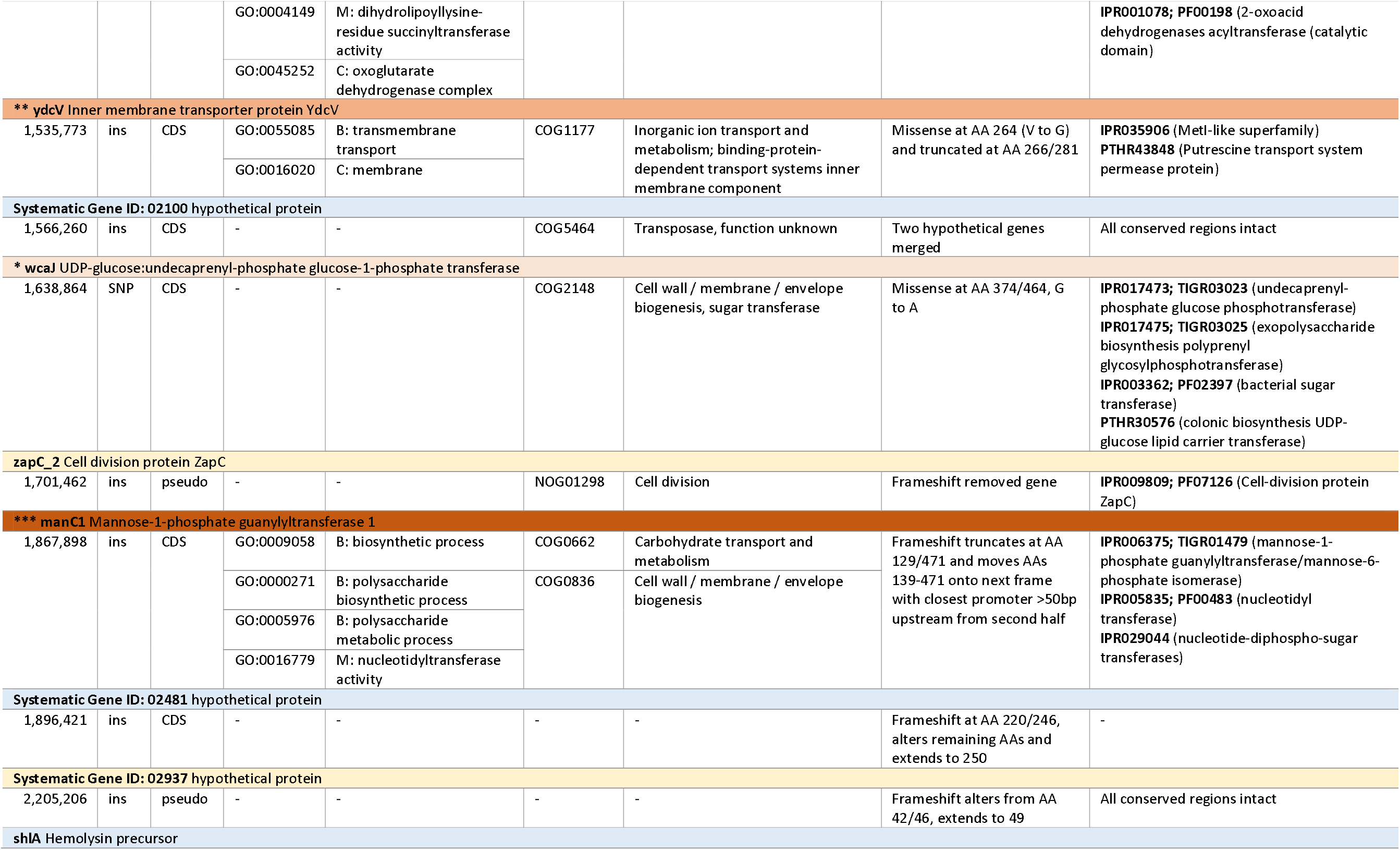
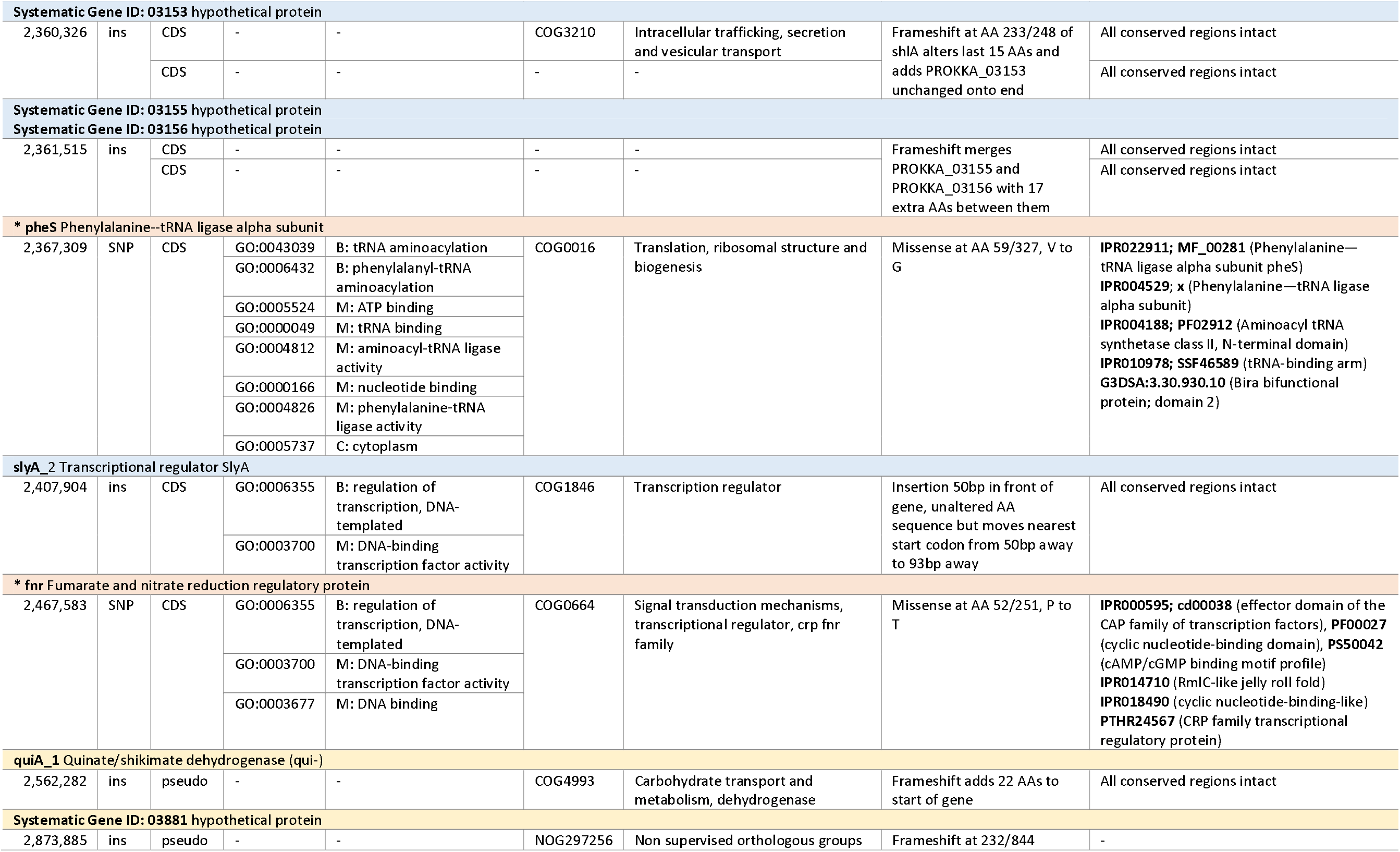
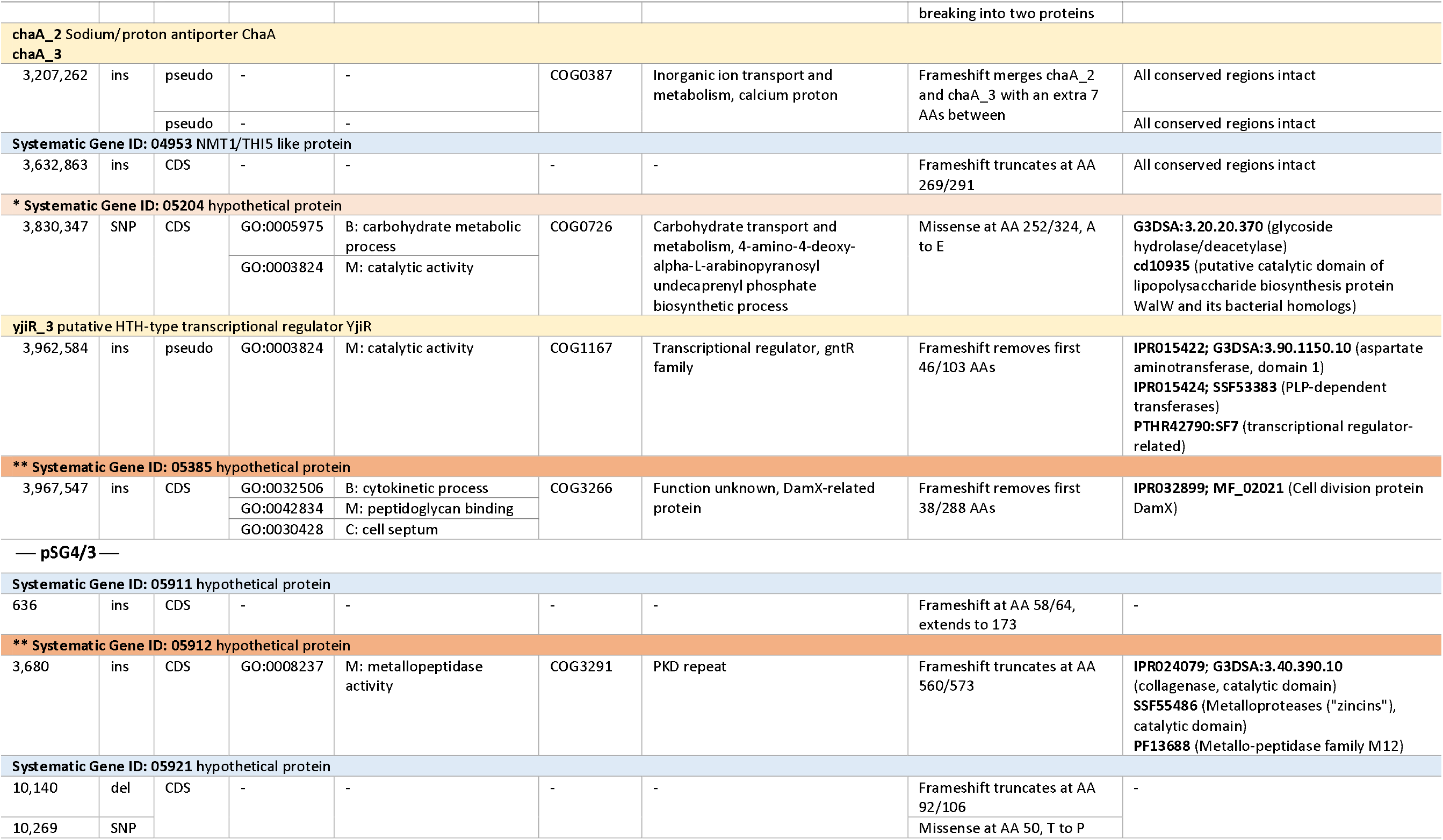
Snippy output with all insertions, deletions and single-nucleotide polymorphisms in *Sodalis glossinidius* strain SgGmmC1*, in comparison with comparator strain SgGmmB4. Gene names and products were annotated using PROKKA; conserved orthologous groups were assigned using the STRING database; gene ontology and conserved regions were assigned using the Interpro database. Key: *** - Tier 1 pseudogenisation (more than two-thirds of the gene as annotated in *Sg*GmmB4 is removed); ** - Tier 2 pseudogenisation (disruption of a conserved region by truncation); * - Tier 3 pseudogenisation (disruption of a conserved region by a missense mutation). CDS – Coding DNA sequence. Pseudo – pseudogene. Ins – insertion. Del – deletion. Com – complex polymorphism. SNP – single nucleotide polymorphism. AA – amino acid. B – biological process. M – molecular process. C – cellular process. Yellow – pseudogenes; blue – CDS; orange – potential novel pseudogenes, shaded by tier.

Raw Nanopore and MiSeq reads are available from the European Nucleotide Archive under Project PRJEB32321 (Read accessions: ERX3321202 and ERX3321201 respectively). A full GBK formatted annotation of SgGmmC1* has been deposited in Figshare at https://doi.org/10.17866/rd.salford.8052437.v1.

## DISCUSSION

The pseudogenisation and deletion of genes described support the hypothesis of symbiont genome size reduction through pseudogenisation and subsequent deletion of genes unnecessary to survival in a stable host environment. In the case of SgGmmC1*, 10 years of passaging in a minimal media with carefully controlled temperature and atmosphere has resulted in inactivation or deletion of genes unnecessary to this environment. Within the 17 kbp deletion most coding genes and pseudogenes were involved in transmembrane transport of organic and inorganic compounds, mainly of carbohydrates, or were membrane-bound transferases. In tsetse, as in other organisms, thiamine is an essential cofactor for amino acid and carbohydrate metabolism that is not present in blood meals. The tsetse fly lacks the capacity for thiamine biosynthesis but carries genes for thiamine transporters (International Glossina Genome Initiative, 2012). *W. glossinidia* and *S. glossinidius* contribute metabolically complementary cofactors to produce an intact thiamine biosynthesis pathway: *W. glossinidia* has the capacity for B vitamin biosynthesis and the synthesis of thiamine monophosphate from which *S. glossinidius* also benefits (Belda *et al*., 2010; Snyder *et al*., 2012; Hall *et al*., 2019).

Serially passaging *S. glossinidius* in thiamine-rich media *in vitro*, with the repeated population bottlenecks caused by passaging, has resulted in the deletion of the *thiM* gene, a putative hydoxyethylthiazole kinase that is an integral part of the thiamin salvage II pathway that allows *S. glossinidius* to synthesise thiazole phosphate carboxylate (THZ-P; Belda *et al* 2010). ThiM had been retained by the original *S. glossinidius* genome despite the loss of other cofactors in the thiamine-production pathway. That *thiM* is deleted in the experimentally evolved *S. glossinidius* strain SgCmmC1*, in the absence of *W. glossinidia* but in the presence of thiamine-rich media, supports the hypothesis that *thiM* is maintained in *S. glossinidius* despite ongoing genome degradation due to positive selective pressure, likely due to thiamine biosynthesis being essential for tsetse survival and achieved through co-operation with the wider microbiome (Hall, 2019).

*S. glossinidius* also maintains a functional acylated homoserine lactone (AHL)-based quorum sensing system that putatively modulates gene expression according to density that is hypothesised to aid coordination between symbiotic bacteria during host tissue invasion (Pontes *et al*, 2008; Renoz *et al* 2023). SgGmmB4 contains two functional copies of esaR, which encodes an ohlL-responsive transcriptional regulator, with the deleted copy in SgGmmC2* being proximal to *thiM*. It is therefore likely that the evolved strain maintains quorum sensing functions due to the presence of the functional copy, however whether density-dependent gene expression has been affected will require further study.

In SgGmmC1* multiple sulfur-associated transmembrane proteins were deleted; sulfur is used in the biosynthesis and modification of sulfur-containing amino acids, methionine and cysteine (Mbaye *et al*, 2019). Methionine and cysteine are both produced by *W. glossinidia* for the tsetse host (Bing, *et al*, 2017) and the degradation of their production pathways in *S. glossinidius* have been predicted as it moves towards a symbiotic lifestyle (Belda, *et al*, 2010). Assuming the deleted sulfur and sulfite transmembrane proteins facilitated sulfur compound uptake, their deletion adds evidence to suggest degradation of methionine and cysteine biosynthesis is occurring.

Carbohydrate transmembrane transporters were the most common group of genes containing small mutations between the evolved strain and the ancestor strain from which it was derived, including those involved in transport of proline, mannose, and a GlcNAc repressor. Further gene functions potentially impacted by variation included protein modification, DNA transcription and regulation, membrane production, two phage lytic phase associated genes, energy production and haemolysis (Table 2). One noteworthy pseudogenisation identified in SgGmmC1* occurred in shlA, a haemolysis precursor (Table 2). During axenic culture it was noted that when grown on Columbia blood agar plates, SgGmmC1* did not exhibit the same level of haemolysis as the original *S. glossinidius* isolate counterpart (data not shown), suggesting phenotypic confirmation of an effect of this pseudogenisation event. This implies either that haemolysis is an important trait in the tsetse host, but not one to those grown outside of its native habitat, or that the conditions for haemolysis gene expression are not met *in vitro*, resulting in a lack of positive selective pressure for maintaining the activity of this gene enabling its deletion.

The serial passaging of *S. glossinidius* imitates the repeated population bottlenecks in natural host-symbiont systems, which are known to contribute to symbiont genome reduction. The *in vitro* passages were repeated every 10-14 days, whereas the lab-reared tsetse life cycle takes more than four weeks. The higher frequency of passaging *in vitro* than transmission of *S. glossinidius* between tsetse in vivo may have increased the bottlenecking effect on genome degradation.

The Black Queen Hypothesis describes the co-dependencies between associated organisms in symbiotic relationships; these co-dependencies are formed by reductive genomic evolution driven by repeated population bottlenecks in a restricted environment (Morris *et al*., 2012). The large deletion in SgGmmC1*, accompanied by inactivation and pseudogenisation of genes via indels, supports the concept the S. glossinidus is capable of ongoing genome reduction by steps as described previously (Goodhead and Darby, 2015). Future work will address the effect of additional selection pressures more closely replicating those found in experimental systems from isolates from tsetse hosts to examine such processes in real-world systems and thereby examine the trajectory of *S. glossinidius* towards obligate symbiosis.

Long-term experimental evolution of *S. glossinidius* isolates under varying selection pressures, such as increased oxidative stress or nutrient deprivation, may alter the evolutionary trajectory of *S. glossinidius* and elucidate the roles of gene systems and selective pressures thereon. Similarly, long-term passaging of *S. glossinidius* through tsetse exposed to trypanosomes may result in genetic, epigenetic or transcription changes in the produced *S. glossinidius* strain, increasing our understanding of the tripartite interaction between symbiont, host, and parasite. Similarly, studying the ability of the evolved lab strain SgGmmC1* to infect tsetse flies would allow for examination of the role *S. glossinidius* genes have on trypanosome vector competence.

## ACKNOWLEDGEMENTS

The authors acknowledge support from the University of Salford for genomics and computational infrastructure and the Liverpool School of Tropical Medicine for supporting the tsetse colony work.

## AUTHOR CONTRIBUTIONS

*Sodalis glossinidius* isolates were collected by ACD and maintained by LRH. Genome sequencing and analysis was performed by PP and IG. Further analysis and manuscript preparation and review was performed by PP, LRH, ACD and IG.

